# Characterisation of a temperate phage induced from *Alicyclobacillus acidoterrestris* DSM 3922^T^: the *Alicyclobacillus* phage MMB025

**DOI:** 10.1101/2025.06.28.662109

**Authors:** Inês Carvalho Leonardo, Helena Ferreira, Ana Patrícia Quendera, Maria Teresa Barreto Crespo, Carlos São-José, Frédéric Bustos Gaspar

## Abstract

The fruit juice industry has increasingly faced spoilage incidents attributed to *Alicyclobacillus* (ACB) bacteria. These spore-forming bacteria produce off-flavours and odours that compromise product quality and lead to significant food waste. Their resistance to acidic pH and the pasteurisation processes commonly used in the food industry demands the investigation of new preservation strategies. Bacteriophage-related treatments have emerged as a promising alternative, being considered safe, green, and sustainable. In this study, the type strain *A. acidoterrestris* DSM 3922^T^ and a food isolate, *A. acidoterrestris* MMB007, were used as hosts to isolate new ACB-targeting phages.

A phage was isolated from a soil sample in contact with the type strain, purified, and characterised for its genetic and phenotypic features. The isolated phage, named *Alicyclobacillus* phage MMB025, belongs to the *Caudoviricetes* class and exhibits a narrow host range. Also, phage MMB025 was revealed to be stable under different environmental conditions, maintaining its lytic activity across a wide range of pH values (3 to 12) and high temperatures (up to 1 h at 60 °C). The hybrid assembly of short and long reads revealed a phage genome of 105,243 bp with a GC content of 43.25 %. Sequence mapping to the *A. acidoterrestris* DSM 3922 genome indicated phage DNA integration within the bacterial chromosome, disrupting the *sigK* open reading frame. This suggests that MMB025 may have originated from a prophage induction event potentially linked to sporulation regulation. Despite the limitations derived from its likely temperate nature, phage MMB025 remains a valuable source of lysins capable of specifically and effectively eliminating ACB bacteria.

More importantly, this study highlights the potential of phages as an alternative preservation strategy against ACB bacteria. The phage MMB025 characterisation opens venues to develop strategies for mitigating ACB-related spoilage, thereby enhancing the sustainability of food production.

## 1 Introduction

Food loss and waste are major global challenges that impact both food security and environmental sustainability. One of the leading causes of food waste is microbial spoilage, which results from the growth of bacteria, yeasts, and moulds. These microorganisms can compromise the safety and quality of food products. The specific spoilage microorganisms that affect various food categories vary depending on multiple aspects, including the intrinsic properties of the food, the type of packaging used, and the environmental conditions during storage (Karanth et al., 2023; Snyder et al., 2024). In the fruit juice industry, spoilage incidents have increasingly been attributed to bacteria of the *Alicyclobacillus* (ACB) genus. These Gram-positive non-pathogenic bacteria produce off-flavours and odours that are described as medicinal, disinfectant-like, and cheesy, compromising product quality and leading to food waste (Leonardo et al., 2025). ACB are particularly problematic because they are thermoacidophilic bacteria, allowing them to withstand the acidic conditions and pasteurisation processes commonly used in fruit juice production. Moreover, their ability to form resilient endospores makes them a persistent challenge for the industry (Cerny et al., 1984; Deinhard et al., 1987; Sokołowska et al., 2020). Thus, alternative preservation strategies are crucial to mitigate ACB-related spoilage events.

Over the years, a variety of non-thermal disinfection strategies has been explored. Industries typically implement stringent cleaning and sanitation protocols, often involving oxidising agents (e.g., peracetic acid). It is well established that high hydrostatic pressure, sometimes combined with mild heat, or along with the incorporation of essential oils or natural antimicrobials, can help eliminate ACB bacteria (Aneja et al., 2014; Guo et al., 2025; Huertas et al., 2014; Silva et al., 2012). However, with the increasing consumer demand for healthy eating, clean-label products, and sustainable food practices, environmentally friendly preservation methods are becoming increasingly popular. Among these, bacteriophage-related treatments have emerged as a promising approach, aligning well with the principles of green and sustainable food protection technologies. These viruses that exclusively target and infect bacteria can be used to control both food-borne pathogenic and spoilage bacteria, offering great advantages compared to traditional chemical and physical treatments. Bacteriophages (or simply ‘phages’) have high specificity, offering a targeted approach to eliminate harmful bacteria without disrupting beneficial microbial communities. Additionally, phages can be directly applied to food products, as they are biodegradable, leave no toxic residues, and do not alter the organoleptic properties of the products (Połaska and Sokołowska, 2019).

The use of phage-based products in the agri-food sector is regulated by authorities such as the European Food Safety Authority (EFSA) and the United States Food and Drug Administration (FDA). In the United States, phage products have been granted the generally recognised as safe (GRAS) status by the FDA, indicating that they are considered safe under their intended conditions of use, and therefore do not require premarket approval. However, companies may voluntarily submit a GRAS notice for FDA review (Liu, S., et al., 2022). In contrast, the regulatory framework in Europe is more restrictive. EFSA excluded phages from its qualified presumption of safety (QPS) list based on an ambiguous taxonomic position or possession of yet unknown molecular traits, and therefore, requires a specific assessment for each phage for which an application is made (Koutsoumanis et al., 2023).

Several countries have already approved phage-based products for application in food, including the European Union, Switzerland, Israel, Canada, the US, Australia and New Zealand. Most of these products are phage cocktails composed of strictly lytic phages that is, phages that upon host infection can only follow the lytic pathway that culminates in bacteriolysis for virion progeny release (Lee et al., 2023; Moye et al., 2018). For instance, ListShield™ targets *Listeria monocytogenes* in ready-to-eat meat and poultry products, fresh and processed fruits and vegetables, and dairy products, while EcoShield™ combats *Escherichia coli* O157:H7 in meat products, being both approved and used in the US and Canada (Carter et al., 2012; Perera et al., 2015; Potera, 2013). In Europe, PhageGuard Listex™ has been approved by EFSA to be applied in food and food processing environments, eliminating *L. monocytogenes* (Moye et al., 2018). Also, phage-based products such as Biolyse® and BAFASAL® have been approved for agricultural applications, including the control of plant pathogens responsible for salad spoilage, potato soft rot, and mushroom blotch, as well as for eliminating human-pathogenic *Salmonella* spp. in poultry farming (Połaska and Sokołowska, 2019). As an alternative to whole phage or phage cocktails, endolysins, phage derived enzymes capable of degrading bacterial cell walls, can be applied in food products (Lee et al., 2023; Potera, 2013). This alternative is particularly interesting when only temperate phages are available. When these phages engage the lysogenic pathway, generally with genome integration into the bacterial chromosome, host cells are not killed and become resistant to subsequent infections by the same or related phages (immunity to superinfection). This feature, associated with the capacity of temperate phages in mediating horizontal gene transfer, impose some limitations to their use as antibacterials (Hyman, 2019). Until now, the only commercially available endolysin approved in Europe is Staphefekt™, which is used not in the food industry but in dermatological applications to treat *Staphylococcus aureus* infections on human skin (Połaska and Sokołowska, 2019; Totté et al., 2017).

Regarding ACB-specific phages, publicly available information is scarce. In 1977, Sakaki *et al*. characterised a lipid-containing phage isolated from a hot spring that targets an *Alicyclobacillus acidocaldarius* strain. In this earlier investigation, the phage’s growth, chemical composition, and stability under different environmental conditions (e.g., range of pH values, temperatures, and different solvents) were evaluated. However, the temperate or lytic nature of the phage was not assessed (Sakaki et al., 1977). Recently, phage KKP 3916 targeting different ACB strains from three distinct species, *A. acidocaldarius*, *A. acidoterrestris, and A. fastidiosus,* was described (Shymialevich et al., 2023). However, this phage carried several genetic elements indicating a temperate lifestyle, making it not ideal for food biocontrol for the reasons explained above. Thus, the study of ACB-specific phages should be further explored as they can serve as an excellent alternative to eliminate these spoilage bacteria from food products while meeting the clean-label demands of the food industry.

In preliminary studies, *A. acidoterrestris* DSM 3922^T^ and a food isolate, *A. acidoterrestris* MMB007, were evaluated as potential targets for phage isolation from distinct environmental samples, including soil and fruit processing water samples. A phage plaque was successfully isolated when testing a sample of vineyard soil against *A. acidoterrestris* DSM 3922^T^. However, during the purification process, the isolated phage lost its ability to infect this strain while being effective against the food isolate *A. acidoterrestris* MMB007. Considering this, the present study aimed to: (1) characterise the isolated phage, regarding its genetic and morphological features, (2) asses the phage host range within the ACB genus, hence, its specificity, (3) evaluate the phage’s stability across a range of pH values and temperatures, and (4) predict the phage lifestyle, either temperate or strictly lytic. Together, these objectives aim to advance the characterisation of ACB-targeting phages, understanding their potential application in the food industry, and offering an alternative food preservation strategy to effectively eliminate these spoilage bacteria.

## 2 Materials and Methods

### 2.1 Bacteria and growth conditions

The bacterial strains used in this study are listed in Table 1.

**Table 1.** Bacterial strains utilised in this study. Culture collection strains and food industry isolates marked with * were generously supplied by Dr. Abby Snyder and Dr. Katerina Roth from Microbial Food and Spoilage Lab, Cornell CALS, Ithaca, NY, USA.

| Bacterial strain | Origin | Reference |
| --- | --- | --- |
| Culture collection strains |  |  |
| *Alicyclobacillus acidiphilus* DSM 14558<sup>T</sup> | Acidic beverage | DSMZ (Germany) |
| *Alicyclobacillus acidocaldarius* DSM 446<sup>T</sup> | Acid hot spring | DSMZ (Germany) |
| *Alicyclobacillus acidoterrestris* DSM 3922<sup>T</sup> | Soil | DSMZ (Germany) |
| *Alicyclobacillus contaminans* DSM 17975<sup>T</sup> * | Soil | DSMZ (Germany) |
| *Alicyclobacillus cycloheptanicus* DSM 4006<sup>T</sup> | Soil | DSMZ (Germany) |
| *Alicyclobacillus cycloheptanicus* ATCC 49029 * | Soil | ATCC (USA) |
| *Alicyclobacillus fastidiosus* DSM 17978<sup>T</sup> | Apple juice | DSMZ (Germany) |
| *Alicyclobacillus fructus* DSM 112018<sup>T</sup> * | Mixed fruit juice concentrate | DSMZ (Germany) |
| *Alicyclobacillus herbarius* DSM 13609<sup>T</sup> * | Dry flower of *Hibiscus* | DSMZ (Germany) |
| *Alicyclobacillus hesperidum* DSM 11985<sup>T</sup> | Solfataric soils | DSMZ (Germany) |
| *Alicyclobacillus kakegawensis* DSM 17979<sup>T</sup> * | Soil | DSMZ (Germany) |
| *Alicyclobacillus macrosporangiidus* DSM 17980<sup>T</sup> * | Soil | DSMZ (Germany) |
| *Alicyclobacillus mali* DSM 112016<sup>T</sup> * | Pasteurised pear juice | DSMZ (Germany) |
| *Alicyclobacillus pomorum* DSM 14955<sup>T</sup> * | Mixed fruit juice | DSMZ (Germany) |
| *Alicyclobacillus sacchari* DSM 17974<sup>T</sup> * | Liquid sugar | DSMZ (Germany) |
| *Alicyclobacillus suci* DSM 112017<sup>T</sup> * | Apple juice | DSMZ (Germany) |
| *Alicyclobacillus tengchongensis* ATCC BAA2134<sup>T</sup> * | Soil of a hot spring soil | ATCC (USA) |
| *Alicyclobacillus vulcanalis* ATCC BAA915<sup>T</sup> * | Hot spring pool water | ATCC (USA) |
| *Bacillus subtilis* CECT 356 | Unknown | CECT (Spain) |
| *Bacillus cereus* BGSC 6A1 | Unknown | BGSC (USA) |
| *Escherichia coli* ATCC 25922 | Clinical isolate | ATCC (USA) |
| *Geobacillus stearothermophilus* DSM 22<sup>T</sup> | Deteriorated canned food | DSMZ (Germany) |
| *Staphylococcus aureus* ATCC 6538 | Clinical isolate, human lesion | ATCC (USA) |
| Food industry isolates |  |  |
| *Alicyclobacillus acidocaldarius* MMB020 | Tomato paste | This study |
| *Alicyclobacillus acidoterrestris* MMB001 | Passionfruit concentrate | This study |
| *Alicyclobacillus acidoterrestris* MMB002 | Peach juice | This study |
| *Alicyclobacillus acidoterrestris* MMB003 | Peach juice | This study |
| *Alicyclobacillus acidoterrestris* MMB004 | Fruit mix concentrate | This study |
| *Alicyclobacillus acidoterrestris* MMB005 | Pear juice | This study |
| *Alicyclobacillus acidoterrestris* MMB006 | Peach juice | This study |
| *Alicyclobacillus acidoterrestris* MMB007 | Tangerine concentrate | This study |
| *Alicyclobacillus acidoterrestris* MMB008 | Sugar | This study |
| *Alicyclobacillus acidoterrestris* MMB009 | Tangerine concentrate | This study |
| *Alicyclobacillus acidoterrestris* MMB010 | Mango and orange juice | This study |
| *Alicyclobacillus acidoterrestris* MMB011 | Peach juice | This study |
| *Alicyclobacillus acidoterrestris* MMB012 | Peach juice | This study |
| *Alicyclobacillus acidoterrestris* MMB013 | Peach juice | This study |
| *Alicyclobacillus acidoterrestris* MMB021 | Peach juice | This study |
| *Alicyclobacillus acidoterrestris* FSL R14-0029 * | Apple juice | Not yet published |
| *Alicyclobacillus acidoterrestris* FSL R14-0046 * | Apple juice | Not yet published |
| *Alicyclobacillus suci* FSL-R14-0023 * | Apple juice | Not yet published |
| *Alicyclobacillus suci* FSL-R14-0051 * | Apple juice | Not yet published |

ACB strains, both from culture collections and food isolated strains, were routinely grown in BAT agar (BAT broth supplemented with 1.5 % (v/v) agar) (Scharlau, Spain) and incubated at 45 °C. To prepare liquid cultures, fresh colonies were grown in BAT broth (Scharlau, Spain) at 45 °C with shaking at 180 rpm in a New Brunswick Innova® 42R incubator (Germany). Before use, both BAT agar and BAT broth were adjusted to a pH of 4.0 using a 1 M HCl solution.

*Bacillus subtilis* CECT 356, *Bacillus cereus* BGSC 6A1*, Geobacillus stearothermophilus* DSM 22, *Staphylococcus aureus* ATCC 6538 and *Escherichia coli* ATCC 25922 were also included in this study to evaluate phage host specificity beyond the ACB genus. They were all grown in Luria-Bertani broth (LB), prepared by adding 10 g/L tryptone (MilliporeSigma, Switzerland), 5 g/L yeast extract (Thermo Scientific, US), and 10 g/L NaCl (VWR Chemicals, US), or in Luria-Bertani agar (LA) if supplemented with 15 g/L agar. Incubation was performed at 37 °C, except for *G. stearothermophilus* where the temperature was 55 °C, with the addition of a 180 rpm shaking in a New Brunswick Innova® 42R (Germany) for liquid cultures.

### 2.2 Phage isolation and purification

A sample of vineyard soil collected from Quinta do Marquês in Oeiras, Portugal, was processed for phage isolation. To achieve this, 10 mL of BAT broth was added to 5 g of soil and homogenised by vortexing. After 10 min of sedimentation at room temperature (22 °C), the upper phase was filtered using a 0.45 µm syringe PES filter. The supernatant was tested against *A. acidoterrestris* DSM 3922^T^ and *A. acidoterrestris* MMB007 using an overlay agar assay. For this, fresh liquid cultures of each isolate were grown and used to inoculate 5 mL of BAT soft agar (BAT broth supplemented with 0.5 % (w/v) agar) with approximately 10^6^ CFU/mL. A volume of 500 µL of the filtered supernatant was mixed with the inoculated BAT soft agar, which was poured onto a previously solidified 20 mL BAT agar base. After 24 h of incubation at 45 °C, the plates were evaluated for the presence of phage plaques.

The phage initially obtained on a lawn of *A. acidoterrestris* DSM 3922^T^ was purified according to the protocol established by Shymialevich et al., 2023, with a few modifications. Briefly, an individual plaque was recovered using a sterile Pasteur pipette and transferred to tubes containing 1 mL of BAT broth, before being incubated at 45 °C on a digital heating shaking dry bath (300 rpm) for 24 h to allow the viral particles to diffuse from the phage plaque agar plug into the medium. The resulting phage suspension was filtered using a 0.45 µm PES syringe filter and used for a new overlay agar assay by adding 100 µL of the phage suspension to the soft agar inoculated either with *A. acidoterrestris* DSM 3922^T^ or *A. acidoterrestris* MMB007. Five rounds of single-plaque passage purification steps were performed until plates with homogeneous phage plaque morphology were obtained. The resulting phage suspensions were stored at 4 °C.

### 2.3 Phage titration

Whenever needed, phage suspensions were quantified (titrated) by determining the number of plaque-forming units (PFU) on a lawn of host bacteria (Clokie and Kropinski, 2009). To achieve this, phage suspensions were serially diluted in BAT broth, and each dilution was mixed with 100 µL of a fresh culture of the target host *A. acidoterrestris* MMB007 in a 1:1 (v/v) ratio. After a 10 min incubation at room temperature (22 °C), the mixture was used to inoculate 5 mL of BAT soft agar, which was poured onto a previously solidified 20 mL of BAT agar base. Plates were incubated at 45 °C for 24 h, the phage plaques were enumerated, and the phage titres were expressed as PFU per millilitre (PFU/mL).

### 2.4 Phage phenotypic characterisation

#### 2.4.1 Transmission electron microscopy analysis

The morphological features of the isolated phage were examined using transmission electron microscopy (TEM) imaging. To achieve this, a fresh phage suspension was prepared and fixed with a 2 % (v/v) formaldehyde solution in a 1:1 (v/v) mixture. A sample of the target bacteria *A. acidoterrestris* MMB007, which had been in contact with phages at a multiplicity of infection (MOI) of 1 for 30 min at 45 °C, was also prepared and fixed using the same approach. Samples were kept on ice until analysis.

Sample negative staining and imaging was performed at the Electron Microscopy Facility at the Gulbenkian Institute for Molecular Medicine. For this, M100 formvar and carbon-coated grids, which had been glow-discharged at 30 mA for 30 s, were used. A volume of 4 µL of each sample was placed on a piece of parafilm, and the grid was positioned on top for adsorption for 2 min. The grid was then washed in 10 drops of distilled water and negatively stained with one drop of 2 % (w/v) uranyl acetate for 2 min before blotting dry. For imaging, an FEI Tecnai G2 Spirit BioTWIN operating at 120 kV was used, and images acquired with an Olympus-SIS Veleta CCD Camera. Images were captured at magnifications of 5k, 105k, and 160k, with overviews at 16.5k. ImageJ was used to determine the dimensions of the virion, measuring the phage head and tail (Supplementary Figure S1).

#### 2.4.2 Adsorption to host cells

To evaluate phage adsorption to *A. acidoterrestris* MMB007, approximately 10^6^ CFU/mL were infected with a phage suspension to achieve a MOI of approximately 0.1 and incubated at 45 °C. Every 15 min, 1 mL of culture was collected and centrifuged at 6000 *g* for 1 min to sediment the cells along with the adsorbed phages. The supernatant was diluted in BAT broth, and the free (non-adsorbed) PFU were quantified as previously described. The number of non-adsorbed phages determined at time 0 min was regarded as 100 %, and the subsequent time points were compared to this sample. The phage adsorption was conducted in triplicate.

#### 2.4.3 Determination of host range

To determine the host range of the phage, agar overlay assays were performed using several bacterial isolates, including multiple species within the ACB genus and others from different genera, such as *Bacillus* spp., *Geobacillus* spp., *Staphylococcus* spp. and *Escherichia* spp. (*Error! Reference source not found.*). To achieve this, fresh cultures of each bacterial strain or isolate were grown and used to inoculate 5 mL of soft agar (BAT or LB 0.5 % (w/v) agar) with approximately 10^6^ CFU/mL (Hossain et al., 2022). The inoculated soft agar was poured onto the previously solidified 20 mL of base agar (BAT or LA 1.5 % (w/v) agar). After drying, 10 µL of serial dilutions of a phage suspension (approximately 10^9^ PFU/mL) were drop-plated on top of the soft agar. After 24 h of incubation, the plates were examined for the presence or absence of phage plaques.

#### 2.4.4 Thermal and pH stability

Phage stability was evaluated across a broad range of temperatures and pH values to assess its resilience to various environmental conditions. To determine phage temperature stability, a suspension containing approximately 10^7^ PFU/mL was incubated at varying temperatures – 4 °C, 20 °C, 30 °C, 40 °C, 50 °C, 60 °C, 70 °C, 80 °C, and 90 °C – for 1 h. The phage titre was determined after exposure to the distinct temperatures. For conditions where the titre fell below the limit of detection, shorter incubation periods were also tested – 30 s, 1 min, and 5 min. Similar tests were performed to assess the pH stability of the phage. For this, a suspension with approximately 10^7^ PFU/mL maintained in standard BAT broth was mixed 1:2 (v/v) with BAT broth prepared at different pH levels, ranging from 2 to 12. After a 1 h incubation, the phage titre was determined. These assays were conducted in triplicate.

### 2.5 Phage genome characterisation

#### 2.5.1 DNA extraction and sequencing

To obtain a phage suspension with the high titre required to extract high-quality DNA (approximately 10^9^ PFU/mL), 20 webbed plates - plates that contain densely packed confluent phage plaques with only a “web” of bacteria left between them - were produced through overlay agar assays. The webbed plates were flooded with 5 mL of BAT broth and incubated for 24 h at 45 °C while shaking at 80 rpm in a New Brunswick Innova® 42R incubator. The phage suspension was recovered, filtered through a 0.45 µm PES syringe filter, and treated with RNase and DNase at 10 µg/mL each for 30 min at 37 °C to digest exogenous bacterial nucleic acids. The treated supernatant was then centrifuged overnight at 4 °C at 15000 *g*. The supernatant was discarded, and the pellet was resuspended in 500 µL of BAT broth by shaking at room temperature at 80 rpm in a New Brunswick Innova® 42R for 2 h. Phage DNA from the resuspended pellet was extracted using the PureLink™ RNA/DNA Mini Kit (Thermo Fisher Scientific Inc., USA) according to the manufacturer’s protocol. The extracted DNA was quantified using a NanoDrop One^C^ Microvolume UV-Vis Spectrophotometer (ThermoFisher Scientific, USA) and sequenced using both Illumina and Oxford Nanopore technologies, as described below.

For short-read sequencing, a genomic DNA library was prepared using the NEBNext® Ultra™ II Library Prep Kit for Illumina® (New England Biolabs, Ipswich, MA, USA) after DNA fragmentation with a Bioruptor Plus by applying 30 cycles of 30 sec/90 sec ON/OFF. The library was quantified using the NEBNext® Library Quant Kit for Illumina® (New England Biolabs, USA), following the manufacturer’s protocol and diluted in 10 mM Tris-HCl to 2 nM. The MiSeq Reagent V2 300 cycle kit (Illumina, San Diego, CA, USA) was used for paired-end sequencing on the MiSeq platform (Illumina, USA), with phi X 174 DNA added to the library as an internal control for the reaction.

Long-read sequencing was outsourced to MicrobesNG (http://www.microbesng.com). Genomic DNA libraries were prepared using the Oxford Nanopore SQK-LSK109 kit with Native Barcoding EXP-NBD196 (ONT, United Kingdom), employing 400 ng to 500lllng of DNA. The barcoded sample was loaded into a FLO-MIN106 (R.9.4.1) flow cell within a GridION system (ONT, United Kingdom).

#### 2.5.2 Genome assembly and annotation

Before proceeding with phage genome assembly and annotation, the preprocessing of short and long reads was carried out using the US Department of Energy Systems Biology Knowledgebase – Kbase (Arkin et al., 2018). Within this open-source software and data platform, the total raw short reads obtained through Illumina sequencing were quality and length filtered using Trimmomatic (v0.39) with a sliding window of 5, a quality cut-off of Q30, and a length cut-off of 50lllbp (Bolger et al., 2014). The Nextera XT adapters were also removed from the reads. The Oxford Nanopore long reads underwent quality and length filtering using Filtlong (v0.2.1) (https://github.com/rrwick/Filtlong) with a quality cut-off of 90 % and a length cut-off of 1,000lllbp.

Following this, a *de novo* assembly was performed using SPAdes (v4.1.0) with the basic command option “--metaviral”, which runs the metaviralSPAdes pipeline for virus detection. As input data, a hybrid assembly was conducted using the previously pre-processed short and long reads (Bankevich et al., 2012; Wick et al., 2017). PhageTerm (v4.1) was used to predict phage genome termini and packaging mode (Garneau et al., 2017), and the final assembly was manually corrected by mapping the pre-processed short and long reads back to the assembled contig using Geneious Prime (v2025.1.2) (https://www.geneious.com), while adjusting the genome start to the newly determined termini. Pharokka (v1.7.5) and phold (v0.2.0) were used to annotate the curated phage genome (Bouras et al., 2023; Heinzinger et al., 2024; Mirdita et al., 2022; Terzian et al., 2021; van Kempen et al., 2024). Finally, the obtained genome was mapped to the *A. acidoterrestris* DSM 3922^T^ genome (Leonardo et al., 2022), aligned using the progressiveMauve command, ran and visualised in Geneious Prime (v2025.1.2) (https://www.geneious.com).

#### 2.5.3 Genome analysis

HHpred was used in the phage genome to predict protein functional domains from annotated CDSs (Söding et al., 2005). DefenseFinder (v2.0.0), the Resistance Gene Identifier (v6.0.5), and the Virulence Factor Database (VFDB) (v4.0) were used in the phage genome to investigate anti-phage defence systems, antimicrobial resistance genes, and virulence factors, respectively (Alcock et al., 2023; Liu, B., et al., 2022; Néron et al., 2023; Tesson et al., 2022). BACPHLIP (v0.9.6), PhageScope (v1.3) and PhageAI (v1.0.0) were employed to predict phage lifestyle (Hockenberry and Wilke, 2021; Tynecki et al., 2020; Wang et al., 2024). Proksee (v1.3.0) was used for genome visualisation (Wishart et al., 2023). The type strain genome was analysed in PHASTEST (v3.0) to predict phage regions (Wishart et al., 2023). The ViPTree server (v4.0) was used to generate a proteomic tree of viral genome sequences based on genome-wide sequence similarities computed by tBLASTx (Mihara et al., 2016; Nishimura et al., 2017). Finally, the tool taxmyPHAGE was employed to search for closely related phages (Millard et al., 2024).

## 3 Results and Discussion

### 3.1 Phage morphological features

The isolated phage, named Alicyclobacillus phage MMB025 (hereafter designated phage MMB025), considering the informal guidelines provided by Adriaenssens and Rodney Brister, 2017, was characterised based on its morphological features. The phage MMB025 forms clear plaques (diameter 1.3 mm ± 0.3 mm) surrounded by turbid halo zones on an A. acidoterrestris MMB007 lawn (Figure 1).

**Figure 1.**
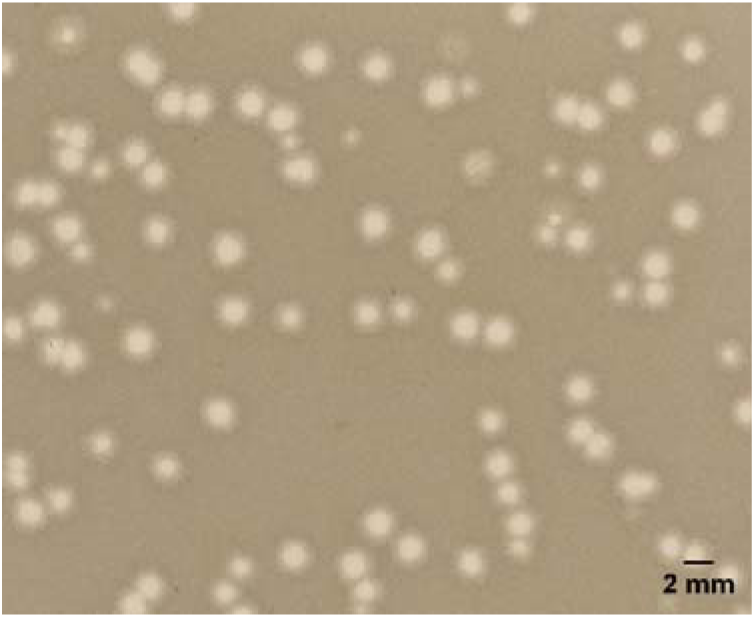
Plaque morphology of *Alicyclobacillus* phage MMB025. Phage plaque morphology on an *Alicyclobacillus acidoterrestris* MMB007 lawn. Scale created in ImageJ.

The analysis of the phage MMB025 through TEM revealed that it consists of an isometric capsid (diameter 65 nm) and a contractile tail (100 × 19 nm) (Figure 2, Supplementary Figure S1). These features suggest that the phage exhibits a myovirus-like morphology belonging to the *Caudoviricetes* class, which encompasses a class of tailed phages whose hosts are bacteria and archaea, accounting for more than 90 % of the total characterised phages (Chevallereau et al., 2022; Zhu et al., 2022).

**Figure 2.**
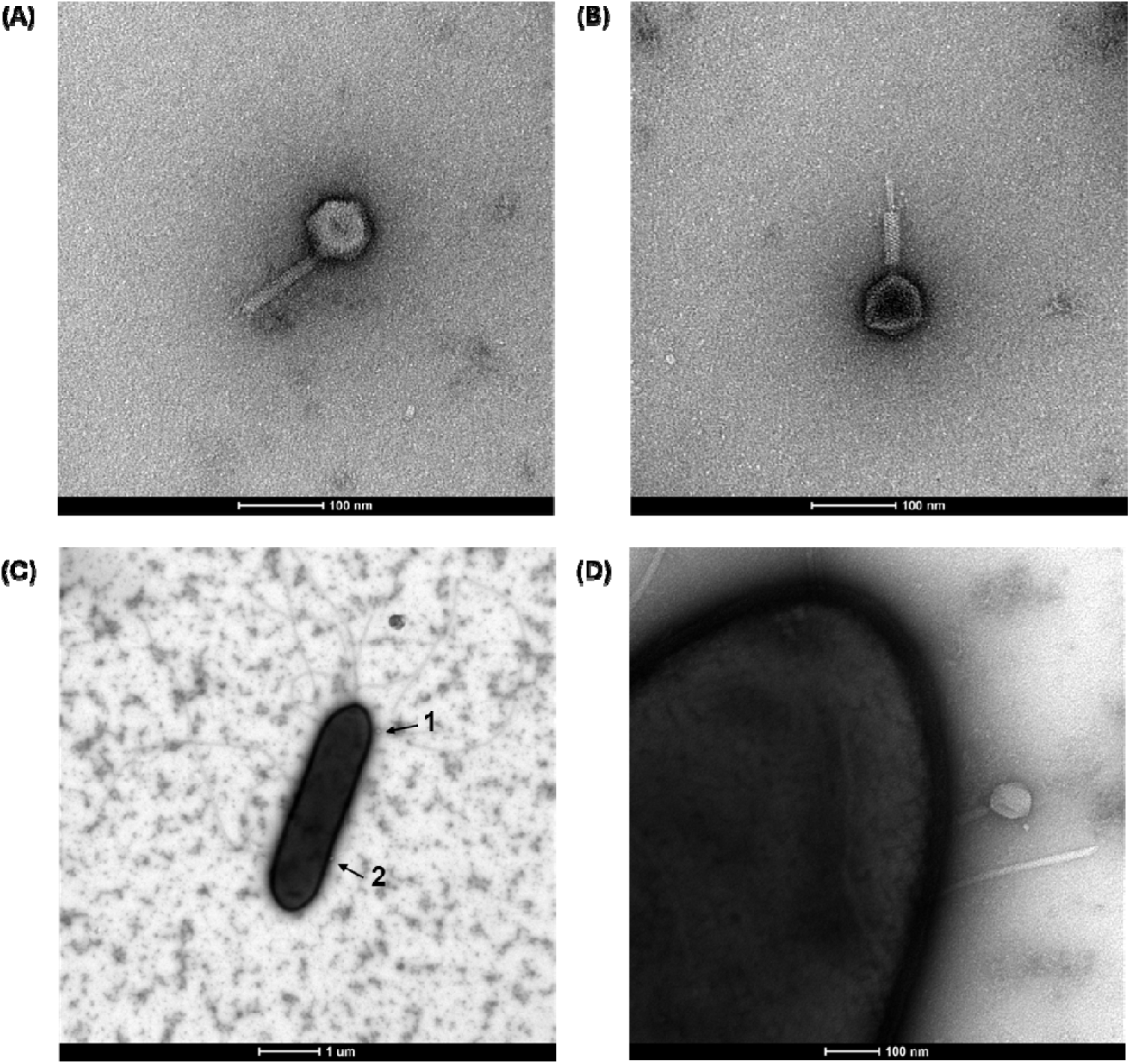
Virions of *Alicyclobacillus* phage MMB025. (A) TEM image of the fixed phage, showcasing the uncontracted tail; (B) TEM image of the fixed phage, featuring the contracted tail; (C) TEM image of A. acidoterrestris MMB007 showing phage MMB025 virions attached to the cell surface (black arrows); (D) Closer TEM image of image C, arrow 1.

### 3.2 Efficiency of phage adsorption to host cells

One of the critical factors influencing the efficiency of phage infection is the rate at which virions adsorb to their bacterial hosts. A high adsorption rate enables more rapid and effective bacterial elimination. As such, adsorption efficiency is a key parameter in evaluating the suitability of a phage for biocontrol applications, including food spoilage control (Abedon, 2023; Moldovan and Wu, 2007). The speed at which phages attach to bacterial cells can be influenced by several factors, including MOI, temperature, pH, and any elements affecting host receptor availability and phage stability. In this study, the adsorption of phage MMB025 to *A. acidoterrestris* MMB007 was evaluated by infecting the host at low cell densities (10^6^ CFU/mL), which already represents a high contamination level in food products. A MOI of approximately 0.1 was selected, and the mixture was incubated at 45 °C. This setup was intended to simulate scenarios unfavourable for phage-host encounter and interaction, while fully supporting the optimal temperature for bacterial host growth, therefore providing a conservative estimate of adsorption performance under suboptimal conditions. The percentage of non-adsorbed phages was then determined at different time points. Under these conditions, phage MMB025 adsorbs to host cells at a slow rate, with approximately 80 % of the phages attaching to bacteria only after 45 min of infection (Figure 3). However, adsorption efficiency may be higher under conditions that favour cell–phage encounters, such as in the presence of higher concentrations of both bacteria and virions. In any case, slow adsorption should not constitute a limitation for the application of the phage in food processing environments, since extended incubation periods during product formulation may compensate for slower adsorption kinetics.

**Figure 3.**
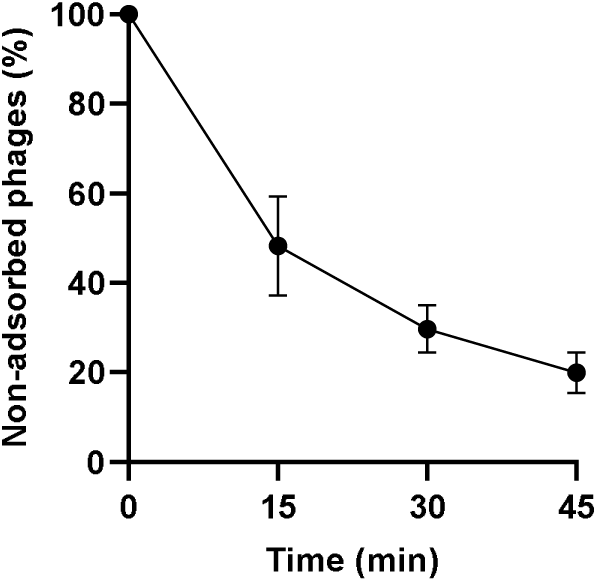
Adsorption rate of *Alicyclobacillus* phage MMB025 to the A. *acidoterrestris* MMB007 host. The presented results are mean values from three independent experiments, with standard deviations indicated by error bars.

### 3.3 Determination of the host range

In addition to phage adsorption rate, the phage host range is a key parameter when evaluating the applicability of phage-based biocontrol strategies within the food industry. To assess the host range of phage MMB025, its lytic activity was tested against different isolates within the ACB genus, including both commercially obtained type strains and isolates recovered from food matrices (Table 1). To further investigate the specificity of this lytic activity beyond the ACB genus, phage MMB025 was tested against bacteria from other genera. These included closely related bacteria such as *Bacillus* spp. and thermophilic bacteria like *G. stearothermophilus*, as well as hygiene indicator bacteria commonly monitored in the food industry, specifically *S. aureus* and *E. coli* (Table 1). These assays revealed the specificity of phage MMB025 for ACB isolates, as no lytic activity was observed against bacteria from different genera (Table 1). The phage MMB025 exhibited a narrow host range, only inhibiting the growth of a limited subset of the tested ACB isolates. Specifically, it inhibited the growth of isolates belonging to only two species: *A. acidoterrestris* (7 isolates), the species most closely correlated with food spoilage events, and *A. acidocaldarius* (1 isolate).

**Table 1.** Lytic activity of *Alicyclobacillus* phage MMB025 against other bacteria. ++: Clear phage plaques were formed; +: Turbid phage plaques were formed; -: No phage plaques were formed.

| Bacterial strain | Phage lytic activity |
| --- | --- |
| *A. acidiphilus* DSM 14558<sup>T</sup> | - |
| *A. acidocaldarius* DSM 446<sup>T</sup> | + |
| *A. acidocaldarius* MMB020 | - |
| *A. acidoterrestris* DSM 3922<sup>T</sup> | - |
| *A. acidoterrestris* MMB001 | - |
| *A. acidoterrestris* MMB002 | ++ |
| *A. acidoterrestris* MMB003 | ++ |
| *A. acidoterrestris* MMB004 | - |
| *A. acidoterrestris* MMB005 | - |
| *A. acidoterrestris* MMB006 | ++ |
| *A. acidoterrestris* MMB007 | ++ |
| *A. acidoterrestris* MMB008 | - |
| *A. acidoterrestris* MMB009 | - |
| *A. acidoterrestris* MMB010 | - |
| *A. acidoterrestris* MMB011 | ++ |
| *A. acidoterrestris* MMB012 | + |
| *A. acidoterrestris* MMB013 | ++ |
| *A. acidoterrestris* MMB021 | - |
| *A. acidoterrestris* FSL R14-0029 | - |
| *A. acidoterrestris* FSL R14-0046 | - |
| *A. contaminans* DSM 17975<sup>T</sup> | - |
| *A. cycloheptanicus* DSM 4006<sup>T</sup> | - |
| *A. cycloheptanicus* ATCC 49029 | - |
| *A. fastidiosus* DSM 17978<sup>T</sup> | - |
| *A. fructus* DSM 112018<sup>T</sup> | - |
| *A. herbarius* DSM 13609<sup>T</sup> | - |
| *A. hesperidum* DSM 11985<sup>T</sup> | - |
| *A. kakegawensis* DSM 17979<sup>T</sup> | - |
| *A. macrosporangiidus* DSM 17980<sup>T</sup> | - |
| *A. mali* DSM 112016<sup>T</sup> | - |
| *A. pomorum* DSM 14955<sup>T</sup> | - |
| *A. sacchari* DSM 17974<sup>T</sup> | - |
| *A. suci* DSM 112017<sup>T</sup> | - |
| *A. suci* FSL R14-0023 | - |
| *A. suci* FSL R14-0051 | - |
| *A. tengchongensis* ATCC BAA2134<sup>T</sup> | - |
| *A. vulcanalis* ATCC BAA915<sup>T</sup> | - |
| *B. subtilis* CECT 356 | - |
| *B. cereus* BGSC 6A1 | - |
| *E. coli* ATCC 25922 | - |
| *G.s stearothermophilus* DSM 22<sup>T</sup> | - |
| *S. aureus* ATCC 6538 | - |

### 3.4 The influence of pH and temperature on phage activity

The assessment of phage stability under various environmental conditions is crucial not only for predicting phage behaviour in the environment but also for evaluating potential industrial applications. Accordingly, the infectious stability of the phage MMB025 was tested by exposing it to different temperatures and pH levels (Figure 4). The phage maintained its titre when subjected to a wide range of temperatures for 1 h, withstanding up to 60 °C with at most approximately a 1 log reduction at this temperature (Figure 4.A). For the higher temperatures of 70 °C, 80 °C, and 90 °C, reduced exposure times were tested, since 1 h exposures resulted in values below the limit of detection (LOD) of the method (Figure 4.B). The phage MMB025 demonstrated a good stability at 70 °C for 30 s and 1 min, with some virions still resisting a 5 min exposure, albeit with higher variations and a significant reduction of at least 5 log (Figure 4.B). At 80 °C and 90 °C, the titre of phage MMB025 significantly decreased compared to the positive control, maintained at 4 °C, even with exposure times as brief as 30 s (Figure 4.B). When exposed to a wide range of pH levels, the phage MMB025 retained its activity at pH values of 3 and above (Figure 4.C). The stability of phage MMB025 under a wide spectrum of pH and temperature conditions—including the acidic environments typical of the fruit juice industry and certain thermal processing temperatures—underscores its potential for application in food products.

**Figure 4.**
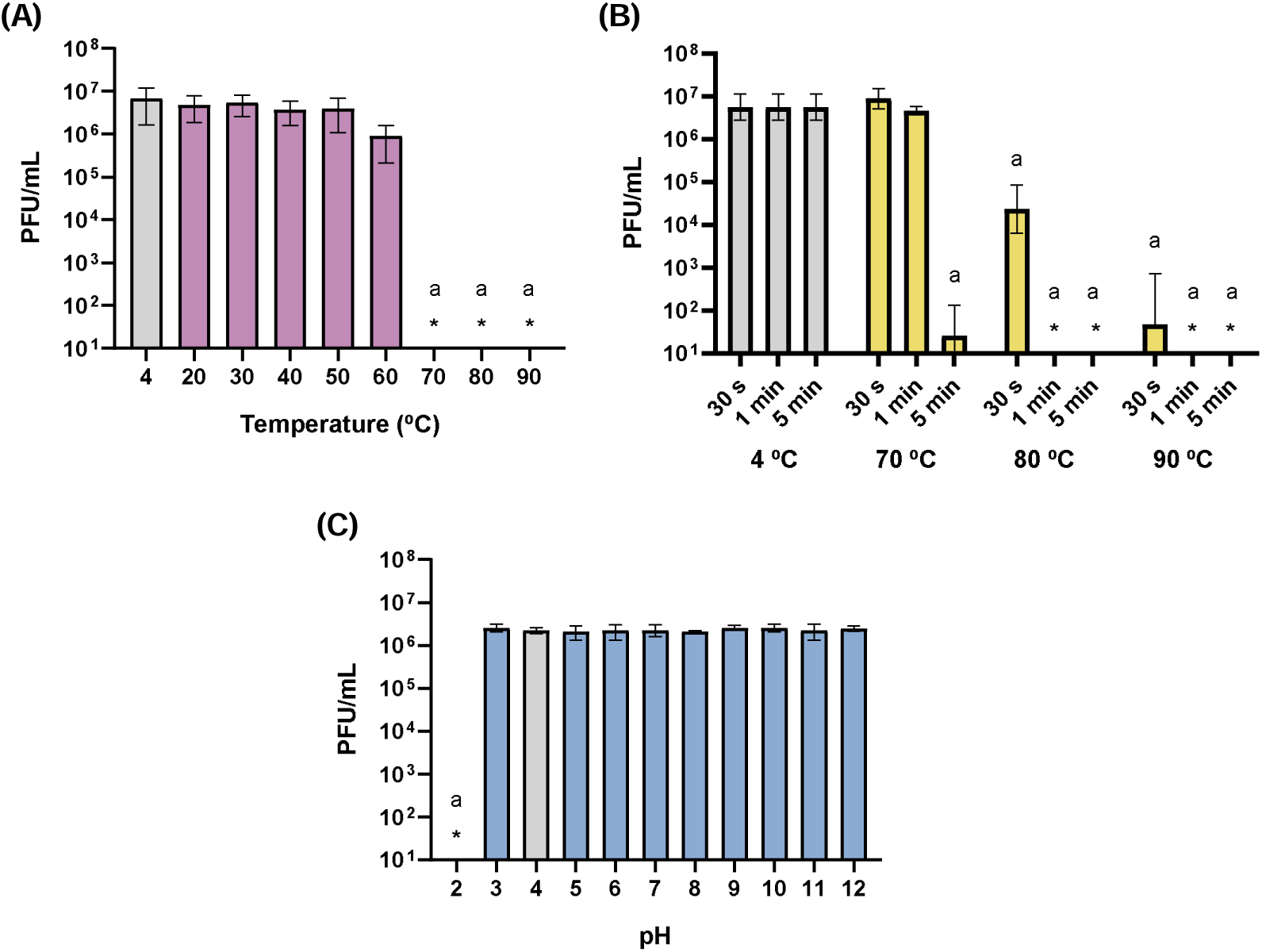
Stability of *Alicyclobacillus* phage MMB025 at different temperatures and pH levels. Stability assessed by titrating phage suspensions following exposure to different conditions: (A) Phage titres after 1 hour at the indicated temperatures; (B) Phage titres after short periods (30 seconds, 1 minute, and 5 minutes) at the highest temperatures (4 °C used as control); and (C) Phage titres after 1 hour at the indicated pH values. Results represent the mean of three independent experiments, with error bars indicating standard deviations. ‘*’ indicates values below the limit of detection of the method. An ordinary one-way ANOVA – for (A) and (C) – or two-way ANOVA – for (B) - was performed for each condition compared to the control (grey bars), which represent the storage conditions of the phage stock solution used in the assay. Conditions with *p*-values < 0.05 are marked with ‘a’.

### 3.5 Phage genomic analysis

The complete genome of the phage MMB025 was sequenced using both short- and long-read sequencing strategies. Typically, only short reads are used to assemble phage genomes; however, hybrid assemblies are known to enable a more robust and reliable analysis and to enhance the confidence in the obtained sequence. Long reads contribute to defining the genome structure, while short reads facilitate a more detailed assembly and individual nucleotide validation (Elek et al., 2023; Necel et al., 2020; Nordstrom et al., 2022; Rozwalak et al., 2024; Zaki et al., 2023).

The hybrid metaviralSPAdes assembled sequence was analysed by PhageTerm, which predicted that the phage has a headful packaging mechanism (Supplementary Table S1). This indicates that rather than packaging a fixed-length and fixed-termini DNA molecule into its capsid, the phage continues to package the DNA until the capsid is full, resulting in terminally redundant genomes, with the packaging process beginning with a cut at a specific starting point (*pac* site) of the phage DNA concatemer. This starting point was identified by PhageTerm and set as the beginning of the genome sequence. After the initial cut at the *pac* site, the DNA is packaged and cut again when the phage head is full, with no predetermined cuts upon packaging terminations, leading to variability in the genome termini. This variability is reflected in an increase in the coverage of the region following the starting point peak, reflecting the region that is present twice in many phage particles (Garneau et al., 2017; Maier et al., 2024). The expected increase in coverage was observed at the same location for short forward, short reverse, and long reads, confirming the packaging mechanism and starting point predicted by the PhageTerm tool. The genome sequence was subsequently manually curated by mapping short and long reads to resolve ambiguities and ensure confident basecalls. The obtained sequence in the form of linear dsDNA consists of 105,243 bp, with a GC content of 43.25 %.

The curated phage genome was first annotated using Pharokka, which uses PHANOTATE to predict coding sequences (CDS) and provide a quick initial annotation (Bouras et al., 2023). Phold was applied in tandem, adding more detailed functional predictions (Heinzinger et al., 2024; Mirdita et al., 2022; Terzian et al., 2021; van Kempen et al., 2024). A total of 214 CDS were predicted, with 73 being associated with genes for which functions could be predicted and 141 CDS related to hypothetical proteins (Figure 5). Among the genes with assigned functions, roles related to DNA replication and transcription regulation, genome packaging, virion structure, lysis, and lysogeny were identified. Regarding structural proteins, five proteins related to the head structure and maturation, the head-tail adaptor protein, two portal-associated proteins, nine proteins related to tail structure, and five baseplate-related proteins were annotated, in addition to two unspecified virion proteins. Pharokka and phold identified two DNA polymerases, four DNA methyltransferases, one DNA primase, one replication initiation protein, and one DnaB-like replicative helicase, among other genes coding for key enzymes ensuring DNA replication and integrity. Eight exonucleases and endonucleases, including four His-Asn-His (HNH) endonucleases and one homing endonuclease, known to be highly abundant in bacteriophage and prophage genomes, were also identified. These endonuclease genes have been suggested to play important roles in the phage lifecycle as key components of phage DNA packaging and in facilitating recombination between phage genomes, contributing to the phage evolutionary dynamics, respectively (Barth et al., 2023; Zhang et al., 2017). A small and a large terminase subunit genes were also annotated in the genome, which are involved in phage DNA packaging into the capsid (Suna et al., 2012; Zhao et al., 2013). Three genes coding for putative integrases were also identified – ORF0038 (14,793 bp to 15,683 bp; 891 bp; 96 aa), ORF0097 (50,899 bp to 52,392 bp; 1,494 bp; 497 aa), and ORF0112 (62,276 bp to 63,382 bp; 1,107 bp; 368 aa) - suggesting that phage MMB025 has a temperate nature, as integrases mediate the integration of the phage DNA into the host genome (Groth and Calos, 2004). The three integrase genes were analysed using HHpred to further investigate their functional domains. HHpred predicts protein structure and function by detecting remote homologies through profile-profile comparisons, searching a variety of curated databases, including PDB, SCOP, Pfam, SMART, COGs and CDD (Söding et al., 2005). All three CDSs were confirmed to contain conserved domains characteristic of recombinases. Comparative analysis revealed homology with both major recombinase superfamilies— tyrosine and serine integrases—distinguished by their catalytic residues (Grindley et al., 2006; Zhang et al., 2023). Specifically, one of the integrases (ORF0038) was predicted to belong to the tyrosine integrase family, while the other two were identified as serine integrases. Pharokka and phold also annotated two genes coding for putative endolysins ORF0078 (37,614 bp to 38,147 bp; 534 bp; 177 aa) and ORF0095 (47,982 bp to 50,243 bp; 2,262 bp; 753 aa). Endolysins are involved in bacterial cell-wall disruption through peptidoglycan degradation at the end of the lytic cycle (Fernandes and São-José, 2018). In this case, HHpred was applied to assess the likelihood of each endolysin possessing catalytic domains characteristic of muralytic activity. One of the endolysins (CDS_0078) was identified as a LysM domain-containing protein with higher homology to tail tube initiator proteins (Veesler and Cambillau, 2011). Despite the LysM domain is involved in peptidoglycan recognition no domains associated with peptidoglycan degradation were detected, suggesting that this protein is more likely to serve a structural role rather than functioning as a lytic enzyme (Mesnage et al., 2014). In contrast, HHpred identified two domains commonly associated with bacterial cell wall degradation in the other endolysin (ORF0095), a peptidase and a muramidase domain, suggesting it is probably responsible for peptidoglycan degradation (Oliveira et al., 2013). In addition to these putative endolysins, genes encoding proteins responsible for fatty acid degradation may also be of interest eliminating ACB bacteria. The ω-alicyclic fatty acids are known to be related to ACB resistance to acidic pH and high temperatures and constitute the major component of ACB cellular membranes, varying from 15 to 91 % of the total fatty acid content (Pornpukdeewattana et al., 2020). In this study, one CDS with predicted enoyl-CoA hydratase/carnithine racemase-like function (ORF0091) was identified in the phage MMB025 genome. This dual annotation reflects sequence homology to and between both enzyme families, which are typically associated with fatty acid metabolism and degradation (Agnihotri and Liu, 2003; Bernal et al., 2007). Further structure and function prediction using HHpred revealed that the 168-amino acid sequence shares similarity with domains found in cysteine peptidases and phospholipases. While this complicates the functional assignment, it suggests a potential role in membrane disruption, possibly contributing to the phage’s ability to infect or lyse ACB bacteria.

**Figure 5.**
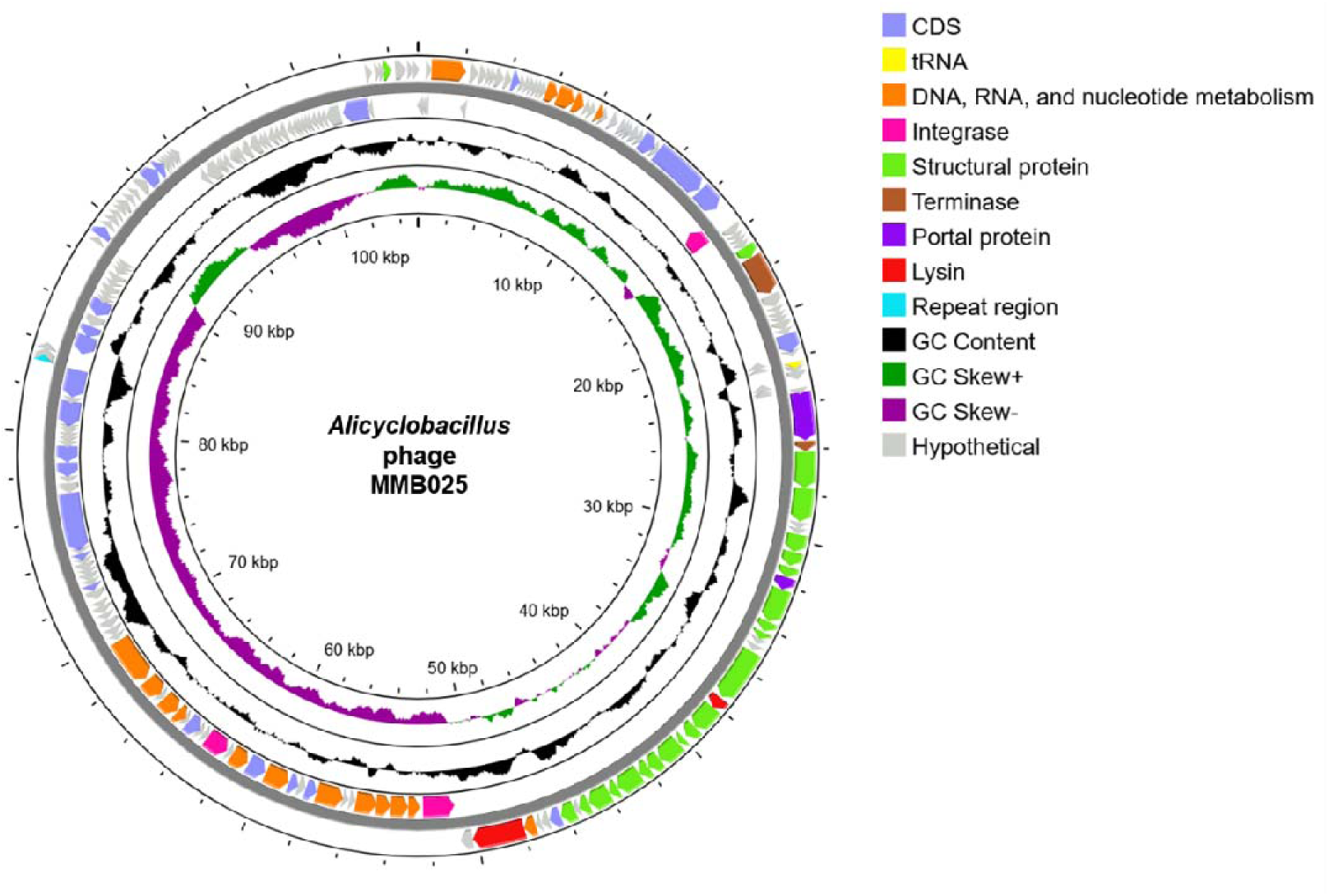
*Alicyclobacillus* phage MMB025 genome. Image adapted from Proksee (Grant et al., 2023). From the outside to the inside: coding sequence (CDS) regions identified on the forward strand, CDSs identified on the reverse strand, GC content, and GC skew. Genetic features and CDS functional groups are highlighted according to the colour code.

The presence of defence systems, antibiotic resistance genes, and virulence factors in the phage genome was also explored using the DefenseFinder programme, the Resistance Gene Identifier application, and the Virulence Factor Database (VFDB) (Alcock et al., 2023; Liu, B., et al., 2022; Néron et al., 2023; Tesson et al., 2022). None of these tools identified anti-phage systems, antibiotic-resistant genes, or virulence factors, respectively. Proksee was used to visualise the phage genome, highlighting the multiple functional groups and regions annotated, namely DNA, RNA and nucleotide metabolism-related CDS, integrases, structural proteins, terminase subunits, portal-associated proteins, and lysis functions (Figure 5) (Grant et al., 2023).

Additionally, the genomic sequence of phage MMB025 was analysed using the viral proteomic tree server VipTree to predict its taxonomy (Nishimura et al., 2017). The analysis of phage MMB025, alongside phages infecting closely related hosts within the phylum Bacillota, determined that it belongs to the Caudoviricetes class (Figure 6), although its specific family could not be identified. The results from VipTree indicate that this phage is well-separated from the classified Herelleviridae family and is more closely related to phages from unclassified families. The direct comparison between the phage MMB025 and the previously reported ACB phage KKP 3916 through MAFFT alignment revealed only 34.3 % identity, suggesting they are not closely related, despite both having Alicyclobacillus spp. as host bacteria (Shymialevich et al., 2023). Furthermore, no matches were identified using the taxmyPHAGE tool to search for closely related phages, suggesting the classification of the phage MMB025 as a representative of a novel genus and species (Millard et al., 2024).

**Figure 6.**
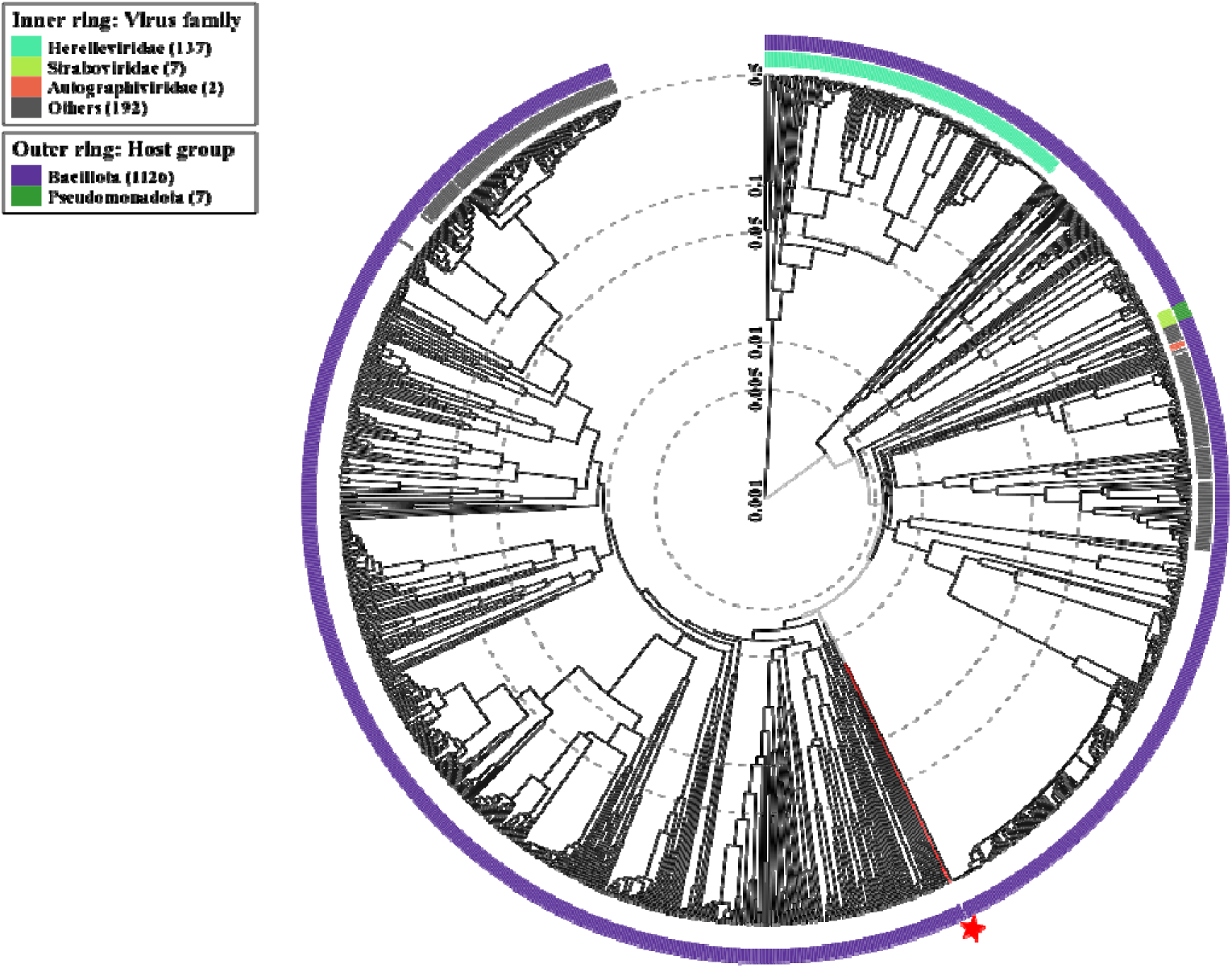
The viral proteomic tree of the *Alicyclobacillus* phage MMB025. The displayed tree was generated using the ViPTree server. It includes the phage MMB025 sequence, all sequences of phages targeting bacteria from the *Bacillota* phylum, and seven phage sequences targeting bacteria from the distant *Pseudomonadota* phylum. Sequence and taxonomic data are based on the Virus-Host DB (Mihara et al., 2016). It was calculated by BIONJ based on the genomic distance matrix and rooted at the midpoint. The tree is presented in a circular view with log-scaled branch lengths. The branch represented by the phage MMB025 is marked in red with a star. The outer rings indicate the host phylum, and the inner rings denote the phage family.

### 3.6 Lifestyle prediction

The lifestyle of a phage is a determining factor in its application as an antibacterial agent. While strictly lytic phages can be directly applied to food products due to their immediate bactericidal activity, temperate phages present challenges, as they can integrate into bacterial genomes and potentially promote phage resistance in target bacteria. In this study, the phage MMB025 was initially isolated from a sample incubated with A. *acidoterrestris* DSM 3922^T^. However, subsequent attempts to propagate the phage in the type strain were unsuccessful, even though it was capable of infecting the food isolate *A. acidoterrestris* MMB007. This may result from the lysogenic nature of *A. acidoterrestris* DSM 3922^T^, which may harbour a prophage that confers immunity against phage MMB025. To assess the potential temperate nature of phage MMB025, its lifestyle was predicted.

As mentioned above, analysis of the phage MMB025 genome revealed genetic elements pointing to a temperate nature. Indeed, BLASTN and MAUVE analyses showed that the MMB025 genome assembled in this study could be fully mapped to the *A. acidoterrestris* DSM 3922^T^, with a pairwise identity of 99.99%, although a rearrangement of genomic sequences was observed (Figure 7). This rearrangement is expected if the recombination site for viral DNA integration in the bacterial chromosome lies roughly at the middle of the phage genome. These findings strongly suggest that MMB025 corresponds to an induced prophage of *A. acidoterrestris* DSM 3922^T^. Interestingly, precise mapping of the bacterial-prophage DNA junctions revealed that the phage is integrated within an ORF encoding a sigma factor (Figure 8). When reconstructed in its uninterrupted form, this ORF shows high sequence identity (> 93 %) to the *sigK* gene found in other ACB species, such as *A. suci* and *A. fructus* (Roth et al., 2021). The integrated phage genome is flanked by inverted repeat (IR) sequences, each followed by a conserved 4-nucleotide motif (TCTC), which likely serves as the core recombination site (RS, Figure 8). This recombination site, located in the position 50,867 bp – 50,892 bp of the phage genome sequence, is immediately upstream of a gene encoding a putative integrase carrying a conserved domain of the SpoIVCA family (ORF0097). This genomic architecture — comprising IRs, a short core recombination motif, and a proximal integrase — is characteristic of site-specific recombination modules commonly found in temperate phages and mobile genetic elements. Although the GAGA motif is unusually short for a recognised recombination site, similar small motifs have been reported in mobile elements of other *Bacillota*. A well-characterised example is the *sigK*-intervening (skin) element in *B. subtilis*, a 48 kb prophage-like region that interrupts the *sigK* gene with a core recombination site composed of only 5 nucleotides (Krogh et al., 1996; Kunkel et al., 1990; Takemaru et al., 1995). Unlike typical prophages, the skin element does not produce phage particles but plays a regulatory role in sporulation. It is precisely excised by the SpoIVCA recombinase during the late stages of sporulation, restoring the functional *sigK* gene and enabling the transcription of late sporulation genes in the mother cell (Molle et al., 2003). This mechanism has also been described in other spore-forming Bacillota, including Clostridium difficile and Clostridium perfringens (Butala and Drago, 2023). Here, we propose for the first time that a similar sporulation-associated regulatory mechanism, mediated by prophage excision, may be present in A. acidoterrestris. The integration of the phage MMB025 within the sigK locus, along with the presence of recombination-associated features, suggests a functional parallel to the skin element. However, unlike the skin element of B. subtilis, which does not give rise to phage particles, the A. acidoterrestris DSM 3922 prophage appears to retain the ability to form infectious phage particles, highlighting a unique dual role in both lysogenic regulation and potential lytic activity.

**Figure 7.**
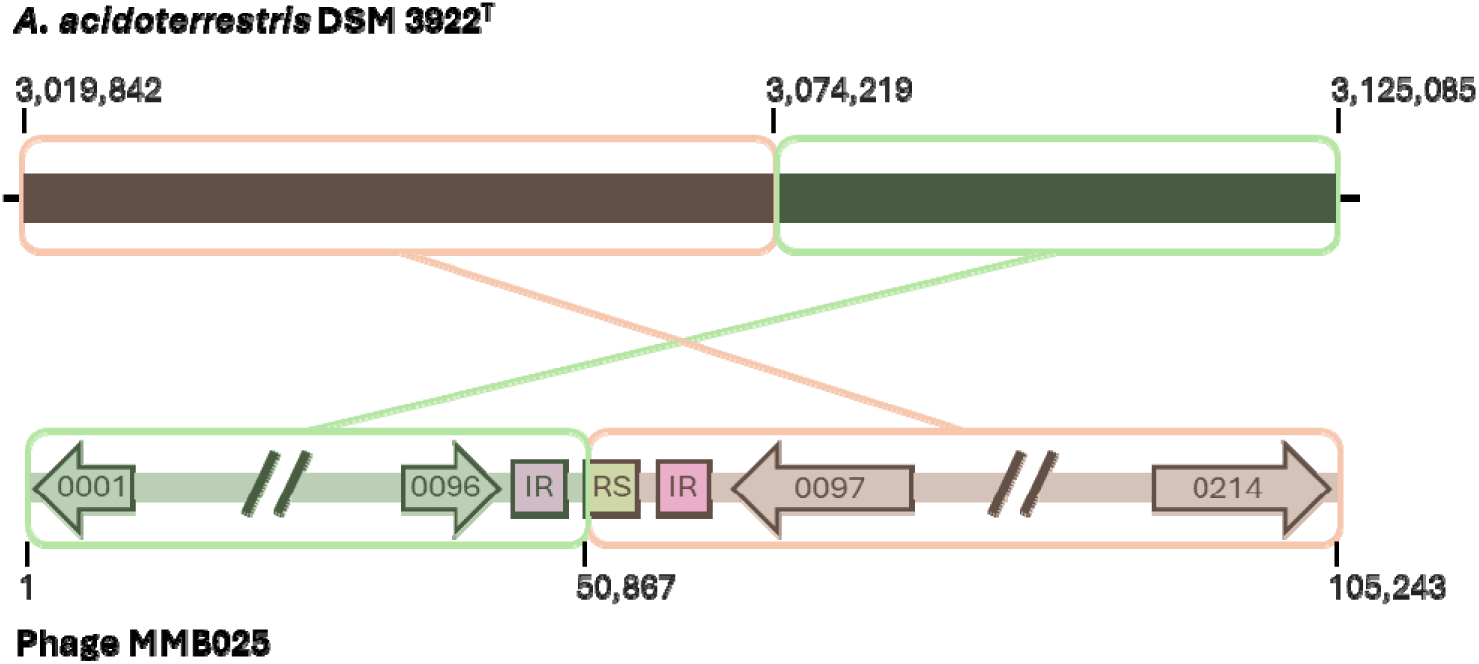
Mapping of the complete sequence of *Alicyclobacillus* phage MMB025 in the *A. acidoterrestris* DSM 3922 genome. The green and red homology boxes highlight the prophage DNA sequence rearrangement, most likely resulting from the site-specific recombination, integration event. Phage MMB025 first and last ORFs, as well as the ones located near the recombination site, are highlighted with arrows. Figures adapted from PHASTEST and MAUVE alignment conducted in Geneious 2025.0.1 (http://www.geneious.com/)(Wishart et al., 2023).

**Figure 8.**
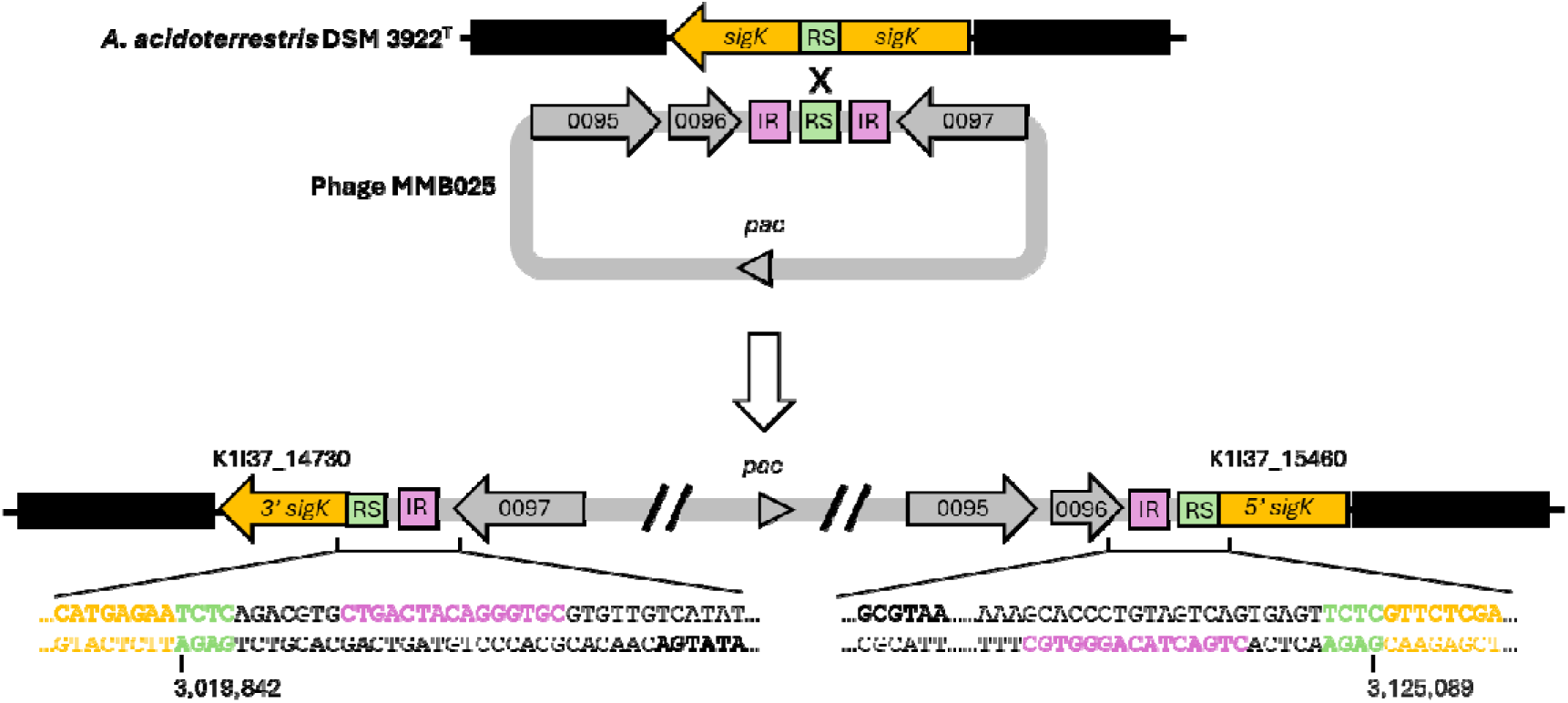
Site-specific recombination between *A. acidoterrestris* DSM 3922 and the *Alicyclobacillus* phage MMB025. The bacterial genome sequence is shown in black, while the phage MMB025 is depicted in grey. Regions identified as inverted repetitions (IR) and recombination sites (RS) are highlighted in pink and green, respectively. The interrupted ORF containing a gene coding for a sigma factor (*sigK*) is highlighted in yellow. The top panel shows the native genomic organisation of the bacterial locus, containing the *sigK* interrupted by RS, and the phage genome containing the pac site, the recombination region with IR and RS, with flanked ORFs highlighted by arrows. The bottom panel displays the resulting recombinant genomic regions, with genomic organisation of the prophage.

Additionally, sequence-based tools can be used to predict a phage lifestyle. In this study, BACPHLIP, PhageScope, and Phage AI tools were tested (Hockenberry and Wilke, 2021; Tynecki et al., 2020; Wang et al., 2024). Considering the phage MMB025 sequence, BACPHLIP revealed that the phage has a 100 % probability of being a temperate phage, while PhageScope suggested a 100 % virulent nature for the phage, and PhageAI predicted a 77.02 % probability of the phage being virulent. The findings of this study underscore the need to improve current bioinformatic tools used to distinguish between lytic and temperate phages based solely on genomic sequences. The observed inconsistencies may originate from the reliance of these tools on structural annotations, which can vary considerably depending on the annotation pipeline employed (Tynecki et al., 2020). To address this, a standardised pipeline with a robust database of phage genome annotations would be necessary to establish a clear, reliable, and consistent distinction between lytic and temperate phages. Therefore, to further elucidate the lifecycle of phage MMB025 and its potential role in sporulation regulation, additional phenotypic and molecular studies are required. Targeted experiments should aim to determine whether the excision of the phage is triggered during late-stage sporulation, thereby restoring a functional *sigK* gene. More importantly, it is crucial to characterise the conditions under which the phage MMB025 transitions between lysogenic and lytic cycles. At this point, and considering its genomic features and origin, phage MMB025 is proposed in this study as a promising source of lysins that effectively target ACB bacteria, rather than as a strictly lytic phage suitable for direct application in food products. To broaden its potential applications, genetic modifications, such as the deletion of the integrase gene, could be explored to convert it into a strictly lytic phage (Sparks, 2020).

## 4 Conclusion

This study presents a comprehensive characterisation of *Alicyclobacillus* phage MMB025, a phage specifically targeting *Alicyclobacillus* spp., a genus strongly associated with food spoilage, particularly in acidic food products. Phage MMB025 was isolated, purified, and characterised for its genetic and phenotypic features, revealing several interesting properties. TEM images revealed their contractile tails, suggesting they belong to the *Caudoviricetes* class. Phage MMB025 exhibits high specificity with a narrow host range while maintaining remarkable stability across a broad spectrum of pH levels and temperatures, including those commonly encountered during fruit juice processing. Genome sequencing and annotation of phage MMB025 revealed that this phage employs a headful packaging mechanism with permuted termini and enabled the identification of more than 70 CDS encoding functional proteins. Among these, a putative endolysin and an enzyme related to lipid metabolism were annotated, which may facilitate bacterial lysis due to peptidoglycan disruption and fatty acid degradation at the end of the phage’s lytic cycle. Phage MMB025 lifestyle prediction was inferred through a comparison between its genome sequence and the *A. acidoterrestris* DSM 3922^T^ genome. This comparison revealed the phage MMB025 to be integrated within the type strain genome, intercepting the ORF coding for the *sigK* gene. This structure is similar to the one described for *B. subtilis* skin element, suggesting phage MMB025 potential role in sporulation regulation. Targeted experiments should aim to determine the conditions under which the phage MMB025 transitions between lysogenic and lytic cycles and to evaluate whether the phage can re-enter the lysogenic state after induction or if excision leads irreversibly to lytic replication and particle formation. Nevertheless, MMB025 can be utilised as a promising source of lysins for addressing spoilage events caused by ACB bacteria in the food industry. Additionally, MMB025 could be genetically modified to expand its host range and convert it into a strictly lytic phage.

Importantly, this study highlights the potential of using phages as an alternative preservation strategy for industrial applications, given that insufficient research has been conducted to specifically address contamination by ACB bacteria in food production settings. Overall, this work reinforces the use of this type of pioneering approach to combat food spoilage through phage-based strategies, which have predominantly been used to tackle food pathogens. It not only expands the current understanding of phage diversity within *Alicyclobacillus* but also lays the groundwork for innovative, GRAS-compliant biocontrol solutions that enhance food quality, reduce economic losses, and thus support sustainable food production.

## 5 Author contributions

**Inês Carvalho Leonardo:** Writing – original draft, Methodology, Investigation, Conceptualisation. **Helena Ferreira:** Methodology, Investigation. **Ana Patrícia Quendera**: Investigation. **Maria Teresa Barreto Crespo:** Writing – review & editing, Supervision, Funding acquisition. **Carlos São-José:** Writing – review & editing, Conceptualisation, Methodology, Funding acquisition. **Frédéric Bustos Gaspar:** Writing – original draft, Writing – review & editing, Supervision, Funding acquisition, Methodology, Conceptualisation.

## 7 Data availability

The data that supports this study is openly available in GenBank of NCBI at https://www.ncbi.nlm.nih.gov under the accession number BankIt2953623 Alicyclobacillus_phage_MMB025 PV686781.

## Acknowledgements

ICL acknowledges Fundação para a Ciência e a Tecnologia (FCT) for the financial support received through PhD scholarship 2020.08210.BD. This work was funded by phages4ACB 2022.04043.PTDC and supported by the Research Units UID/04138: iMed.ULisboa - Research Institute for Medicines, and UID/04462: iNOVA4Health – Programme in Translational Medicine, financially supported by Fundação para a Ciência e Tecnologia / Ministério da Educação, Ciência e Inovação and the Associate Laboratory LS4FUTURE (LA/P/0087/2020). This work was produced with the support of CNCA funded by FCT. The authors also acknowledge the financing of the Agenda TEC4GREEN, n° 02/C05-i01.01/2022. The authors would like to express their gratitude to Ana Laura Vinagre and Michael J. Hall from the Electron Microscopy Facility at the Gulbenkian Institute for Molecular Medicine for performing the TEM imaging, and to Dr. Abby Snyder and Dr. Katerina Roth (Microbial Food Safety and Spoilage Lab, Cornell University, Ithaca, NY, USA) for their contribution to this work.

