## Supplementary Figure S1 for "Characterisation of a temperate phage induced from *Alicyclobacillus acidoterrestris* DSM 3922^T^: the *Alicyclobacillus* phage MMB025"

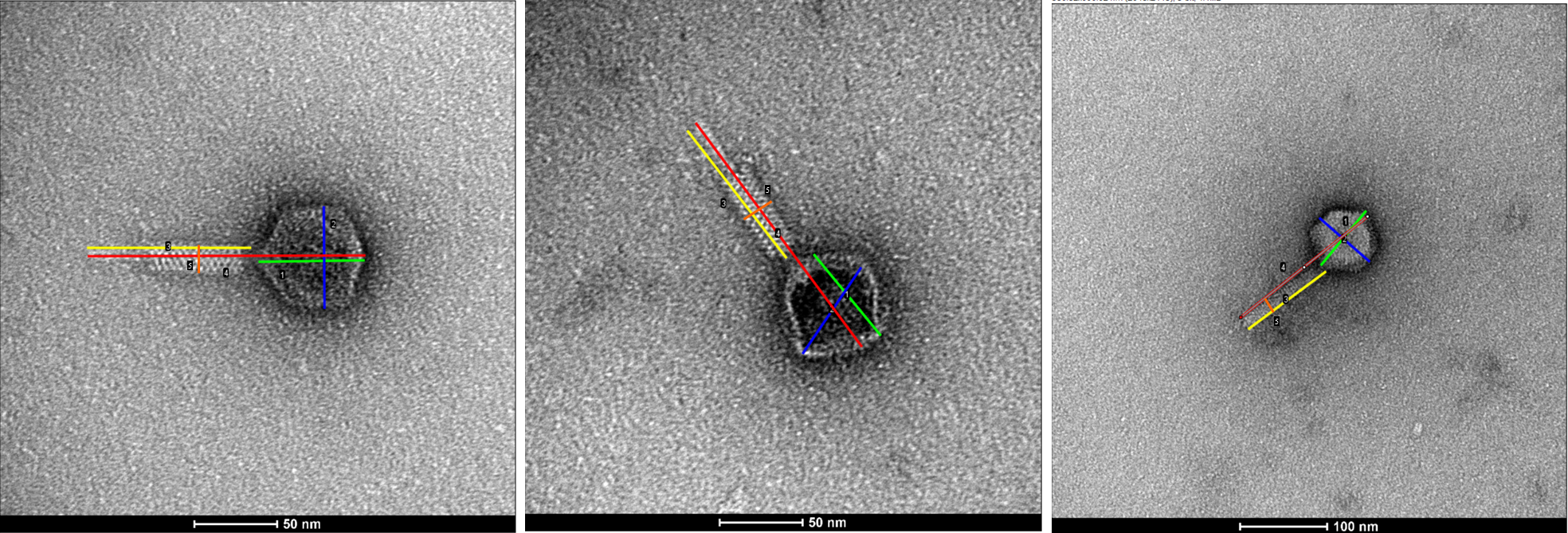

|  | **Figure 1** | **Figure 2** | **Figure 3** | **AVG** | **STDEV** |
| --- | --- | --- | --- | --- | --- |
| **Head length (nm)** | 61.312 | 62.38 | 74.554 | 66 | 6.006455 |
| **Head diameter (nm)** | 59.527 | 61.899 | 73.721 | 65 | 6.208021 |
| **Tail length (nm)** | 95.387 | 96.476 | 107.398 | 100 | 5.423612 |
| **Tail diameter (nm)** | 17.66 | 19.653 | 20.296 | 19 | 1.1222 |
| **Size phage virion (nm)** | 163.096 | 167.045 | 184.724 | 172 | 9.403971 |

**Figure S1 – Measurements of the *Alicyclobacillus* phage MMB025 performed in ImageJ.** Three images of the phage MMB025 were selected, corresponding to the phage with a contracted or extended tail. 1 – head length, green, 2 – head diameter, blue, 3 – tail length, yellow, 4- phage virion size, red, 5 – tail diameter, orange.
